# A Generative Bayesian Approach for Incorporating Biosurveillance Sources into Epidemiological Models

**DOI:** 10.1101/328518

**Authors:** David Manheim

## Abstract

Biosurveillance “systematically collects and analyzes data for the purpose of detecting cases of disease, [and] outbreaks of disease.” (Wagner, Moore and Aryel, 2006) This typically involves using a set of known sources of epidemiological data, instead of opportunistically using the data sources which become available over time. This work attempts to partially remedy that limitation by using an easily adapted generative Bayesian econometric model to allow incorporation of novel data sources. This is done by building a generative model of the information sources, then using Bayesian Markov-chain Monte-Carlo to find the relationships between data and actual caseloads to use in an epidemiological model ^1.^ While the application presented is limited to three data sources for a single disease (influenza), the methodology is potentially widely applicable, and enables rapid incorporation of a variety of sources and source types.

## Introduction

In the past half century, the world has gone from almost no biosurveillance to a complex set of different systems that provide different types of information about different diseases - and the US spends billions of dollars per year on these systems (Wagner, Moore and Aryel, 2006). Our much-improved planning and response depends on these systems, and the impact is overwhelmingly positive. Given new sources and types of data that can pose novel challenges, the proliferation of data and surveillance systems has outpaced our ability to use the various sources of data. (Wojcik et al., 2014)

There are a variety of approaches to integrating data into predictive models, but most require large historical datasets to allow accuracy, and new data sources often require extensive model tuning to add predictive value. Most emerging data sources come without a large corpus of historical data to be used to train them, and they do not allow the incorporation of prior or system knowledge to capture the interrelationships between data sources. To remedy this, the current work built a generative model for information fusion of diverse data types. This uses a multilevel Bayesian “information model” that enables easy expansion to include new influenza biosurveillance data sources, as explained in earlier work (Manheim, 2018). This information model is used to estimate the inputs of a standard compartmental disease model, and this combination of a theory-based SEIR model and a statistical Markov-chain Monte Carlo model has advantages for a variety of applications. (Manheim et al., 2016).

## Model Structure

As mentioned above, this model has two principal components: (1) the epidemiological influenza model, and (2) the information model. Additionally, the model uses a numerical Bayesian method which allows the epidemiological model and the information model to be used together.

The influenza model is a compartmental SEIR model. Individuals in the model are assigned to one of four statuses, or compartments; Susceptible, Exposed, Infected, or Recovered - that is, S, E, I, or R, hence the name. The rate at which people become infected depends on the number of infectious individuals, and so the model uses a system of coupled differential equations to track the movement of so that the rate at which individuals move between these states. The model takes as inputs the characteristics of the virus, the population characteristics, and the initial conditions, and outputs the (unobserved) actual dynamics of the disease over time. This model is deterministic, and excluding minor numerical accuracy issues, the uncertainty in the result comes from the uncertainty in the inputs, and the geographic and population aggregation that occurs.

The data used to inform the model is observations of various types, principally biosurveillance sources. The three sources used are Influenza-like Illness (ILI) and the National Respiratory and the Enteric Virus Surveillance System (NREVSS) data, both collected by the CDC. (Centers for Disease Control and Prevention, 2016) These formed the basis for the initial model, while the third, Google Flu Trends (Ginsberg et al., 2009), was used as an example of how additional data sources could be incorporated into the model^2^. For each source, the information model represents the relationships between the data that is observed, and the ground truth data (or the modeled, unobserved dynamics.)

Combining these models is done using the Stan^®^ modeling platform, (STAN Development Team, 2017) which allows representing the information model and Bayesian priors for the parameters, (Gelman, Lee and Guo, 2015), along with an integrated ODE solver for the epidemiological model (Serban and Hindmarsh, 2003). Using these two models, Stan can perform Hamiltonian Monte Carlo sampling to find both the estimated posterior for the information model, and the estimated outputs of the ODE model.

## Influenza Model

Susceptible-Exposed-Infectious^3^-Recovered (SEIR) models are a standard method for representing influenza and many other similar infectious diseases^4^. The model represented the transition between states; starting with those susceptible, they can be exposed to the disease, and have a latent and developing case, so that they are infected but not yet infectious, at a rate that depends on their contact with infectious individuals. The exposed then become infected at a rate determined by the biology of influenza. After the influenza has run its course, the individual can no longer be infected due to serological immunity, and they may then be considered recovered from a particular strain for multiple years.

This model is a causal representation of how a disease spreads in the population, based on a straightforward logic mode for disease transmission. In the model, the actual process is simplified by representing population groups in aggregate, and tracking the overall rates of contact with infected individuals and the course of the disease. The course of the process is approximated using a set of differential equations that represents the transition between disease states of people in the population over time, as shown in Figure 1: SEIR Model Structure, Inputs, and Differential Equations Governing the Model.

**Figure 1:**
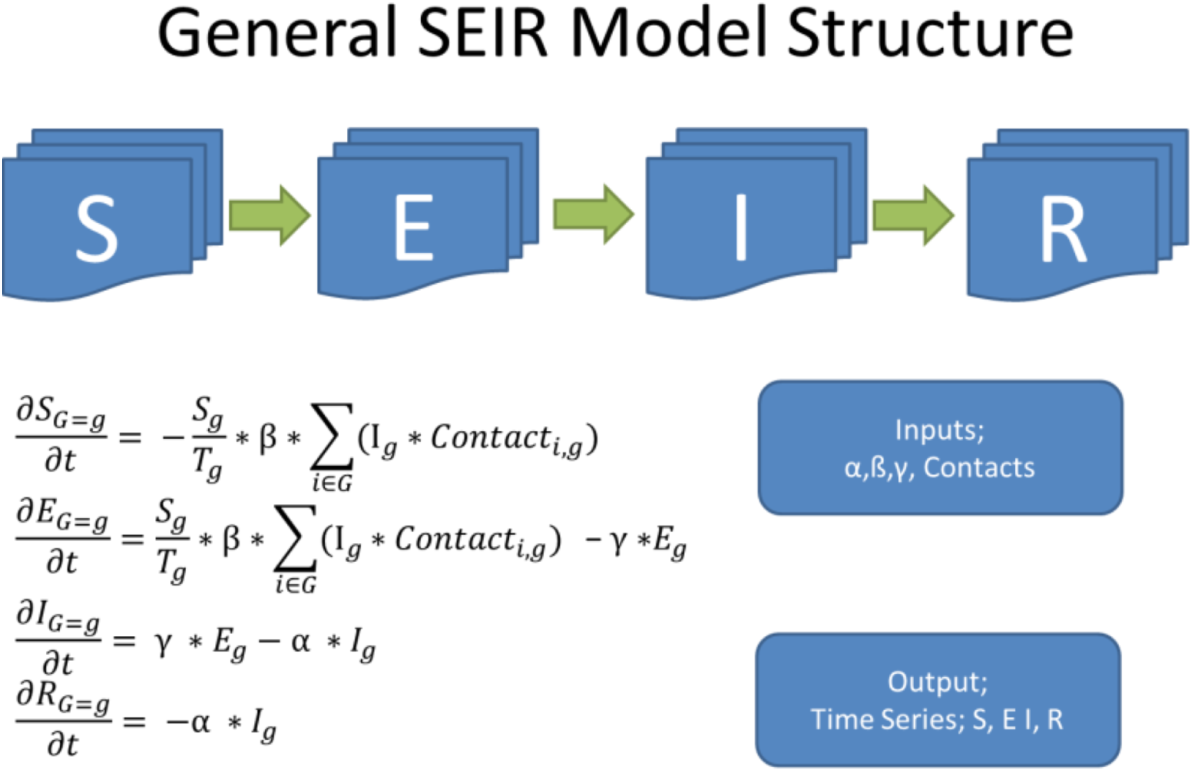
SEIR Model Structure, Inputs, and Differential Equations Governing the Model.

Each of the rates of transition, and several other related parameters, can differ to a greater or lesser extent over time, between different age groups, individual behaviors, and between yearly strains and pandemic influenza. The SEIR model constructed for this project simplifies this into categories representing different age groups,^5^ each of which has some proportion of their population that has each status over time. The differential equations governing the transitionrates are given in the figure, where each group () has 4 compartments (each indicated by their initial) subscripted by population group. The unknown variables α, β, γ represent the probability an infected person infects a contact, the average rate of become infected after exposure, and the rate of recovery, respectively. The contact matrix is the rate at which people in different groups encounter each other.

## Information Sources and Information Model

An information model relates the data available to inputs and outputs of the phenomenological model. The influenza model uses three extant data sources, as outlined earlier. Critically, these relationships form a generative model that not only represents the full relationship between the variables, but does so in a way that matches the causal structure of the information generation process. It can also incorporate varying time structures of data availability. This allows the Bayesian update conditional on the flu data to estimate the relationships in this causal model.

For example, ILI data is collected from a sample of healthcare providers throughout the US, and now includes several thousand providers. It is available approximately 2 weeks after it is collected. Each flu season, the CDC enrolls providers to report weekly data on the total number of patients they see, as well as the number of patients by age group with “fever (temperature of 100°F [37.8°C] or greater) and a cough and/or a sore throat without a known cause other than influenza.” (Centers for Disease Control and Prevention, 2016)

The captured data does not represent the full structure of the information generation process. As an example, some number of people each week do not feel well, due to influenza or for another reason, or visit the doctor for a routine checkup. The total number of such visits at each ILI surveillance site is reported. A subset of such people both exhibit flu symptoms go to a doctor who participates in the flu surveillance system, and the number of such people in each age category will be reported as an influenza like illness. The available data does not capture unobservable features of the system such as the fraction of influenza patients who are symptomatic, nor the fraction of symptomatic patients that go to a doctor.

In order to model this data, we define variables used in the information model which requires a prior estimate, which is then updated based on the data. The structure of the relationship between the auxillary variables and the data for CDC data sources is shown in the illustrative Figure 2 below, and the actual model structure estimated can be seen in the source code for the STAN model. In the diagram, we see the overall causal structure for the information generation process.

**Figure 2:**
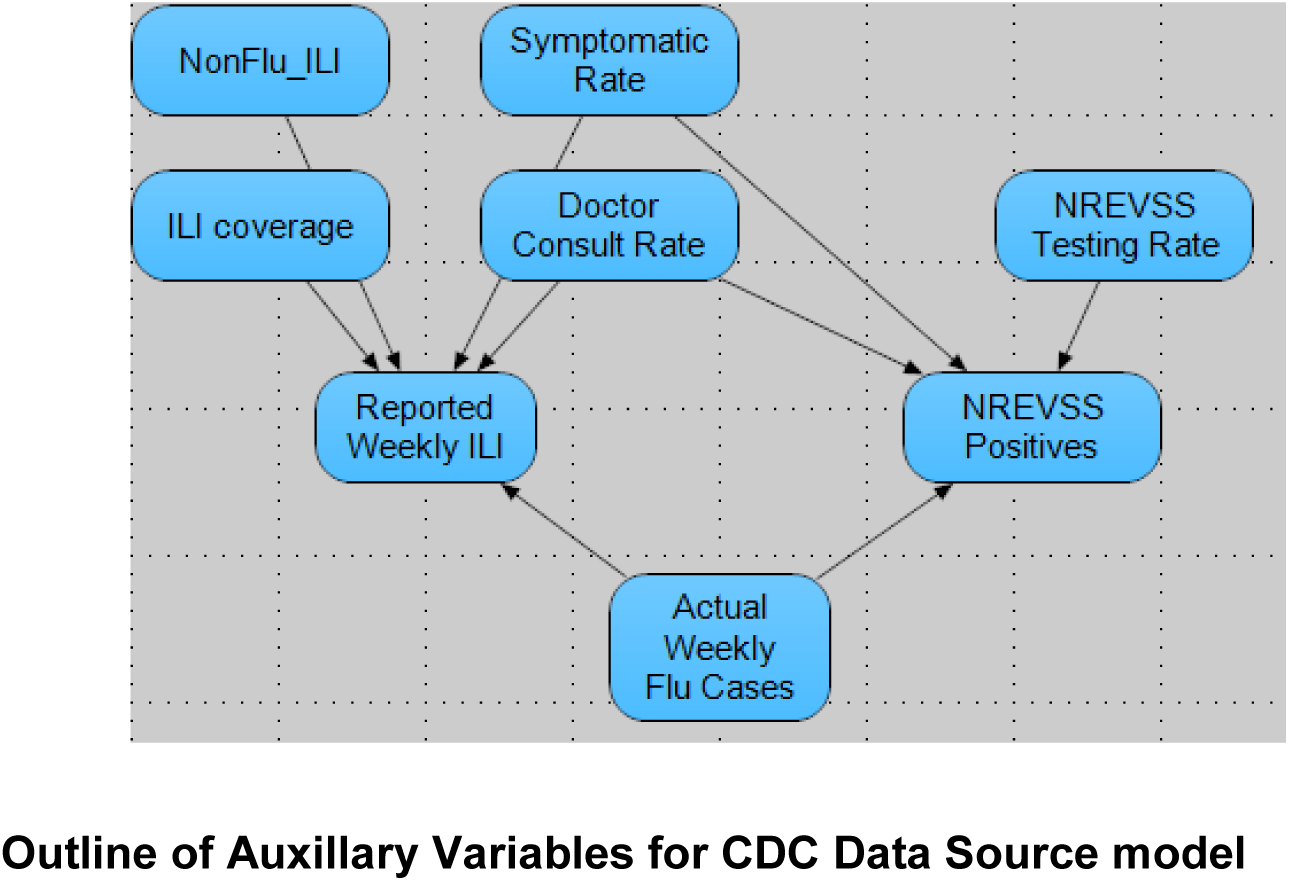
Information model for CDC Data, Illustrative Relationship Diagram.

Outline of Auxillary Variables for CDC Data Source model

- Nonflu ILI is the number of (symptomatic) non-influenza illnesses in the population. The prior used for nonflu-ILI per time period is a gamma distribution, with alpha = 0.01, beta = 0.1.
- The symptomatic rate is the percentage of the infected population that exhibits symptoms. Medical literature suggests that this is around [in equ], so a moderately informative beta distribution prior with alpha=10, beta=30 is used.
- The Consult Rate is the percentage of people with symptomatic flu that go to a doctor, which uses an uninformative prior with mean 10%.
- ILI coverage (ILI_Pop_Pct) is a fixed value defined as the percentage of the total population covered by the ILI network in a given week. It is calculated based on input data, and varies by week because the number of offices/clinics that participate in ILInet varies over the course of the year.
- The NREVSS testing rate is assumed to be the percentage of people who go to a doctor who have a flu test performed; it is simply the total number of NREVSS tests performed divided by the calculated number of average doctor visits per week.

The number of actual weekly flu cases is represented by the number of people in the I compartment in the influenza model. The relationships between the observations and this overall number is represented by the generative information model. Because the sample data has a total number of patients seen, we consider the number of people who consult the given doctors overall, and those who have ILI and go to the doctor for that reason. Of the people who have influenza, only a fraction (that depends on age) are symptomatic. People who go to the doctor do so because they are symptomatic, which can either be due to actual influenza, or due to non-flu ILI.

NREVSS data is virological surveillance data provided by laboratories (now several hundred, including both CDC,s NREVSS system and the WHO laboratories in the US.) These reports are available approximately 4 weeks after the samples tested are collected. If a lab test is performed for a person at a lab which reports to NREVSS, and the symptoms are in fact flu, this data point will appear as a positive case of influenza in the NREVSS data^6^.

## Extending the Information Model

Extending this model to capture additional data sources requires inputting the causal structure of the new data source. This does require significantly more work on the part of the modeler than a typical machine learning model like a neural network, but has the advantage of being able to incorporate any data sources with understood relationships to any of the modeled factors.

In this case, we consider Google Flu Trends data. This source reported the volume of searches for “flu search activity” compared to the long-term baseline. The data implicitly measures a function of influenza-like illness, since searches are presumably in large part performed by individuals who cannot distinguish between flu and similar illnesses. Google Flu Trends data is reported as an index, where the value represents the number of standard deviations above or below the normal volume of influenza related search activity. (Ginsberg et al., 2009) Because the average and standard deviation are not reported, we define them as auxiliary variables, and the Bayesian model finds the appropriate value that best fits the data set given. During non-flu season, which is summer in the United States, the value is near −1, while in a severe flu season it reaches 4 or higher. The model will assume this data is available immediately, to represent the potential of such a system^7^. However, there is much more uncertainty in this estimate; the volume of searches can react to factors other than flu, such as fear of flu as occurred in 2009. We assume that search volume is a function of total number of symptomatic ILI cases, caused by influenza or not. Using these assumptions, we can represent the data in the model.

Importantly, other classes of data relating to the modeled disease can also be used. For example, information about the strains circulating in a given year, for example, can be used to estimate the non-susceptible population proportion for different age groups by extending the information model.

## STAN Model implementation

Standard practice in Bayesian Analysis, as explained by Gelman et al (2014), starts by building a full probability model relating all the observable and unobservable data, as we have sketched in our information model. This should be consistent with the causal relationship between factors, and consistent with the way the data is collected. For instance, if we fail to correctly model inaccuracy in measurements, the model will not represent the relationship between input data and other variables, and the model may fail to converge, or will have high uncertainty in the estimate.

Each of the uncertain inputs in the model, including both the unknown values in the phenomenological model and the auxiliary variables in the information model, are treated as random variables with a weakly informative prior distribution. Conceptually, the posterior values for all of these variables are computed using Bayes, rule. Direct calculation of the posterior is intractable, and modern Bayesian tools use computational estimation methods. Stan uses a set of heuristic approaches to draw an initial sample from the estimated posterior distribution to use for adaptation of the Markov chain, i.e., to reach an area of high posterior probability. These samples are used to initialize the mass matrix and find a useful step size for later sampling. Then, following a warm-up period, Hamiltonian Monte Carlo is used to generate samples from this posterior distribution. The mean values of the posterior are reported in Table 1.

**Table 1.**
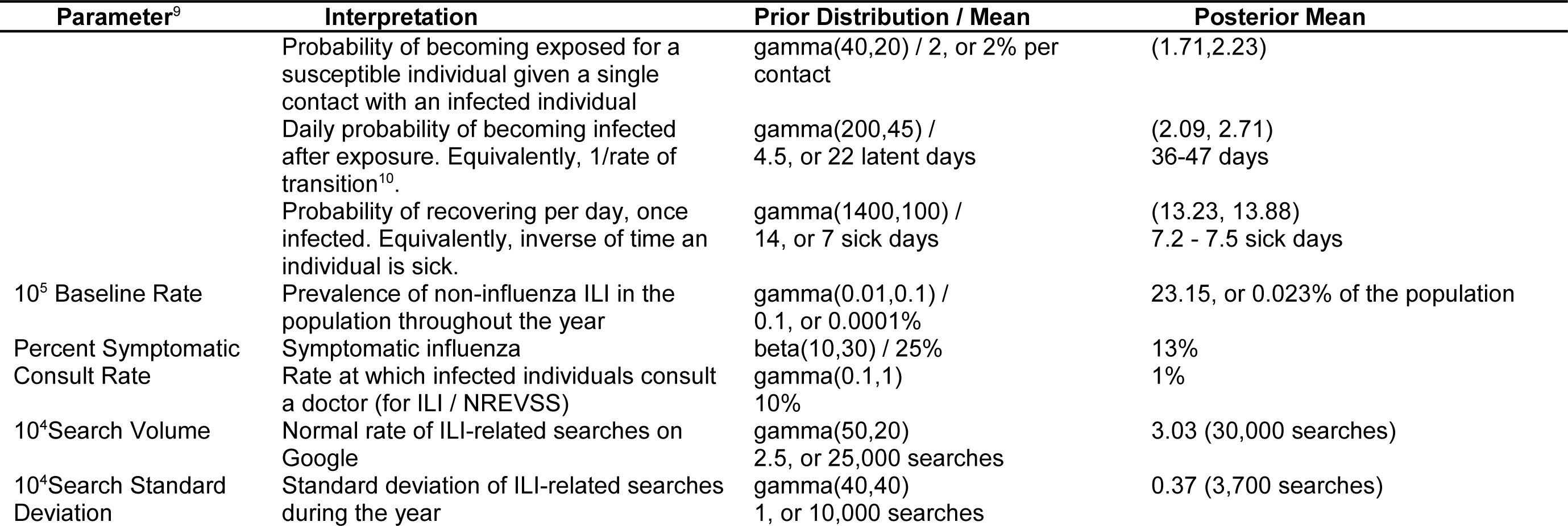
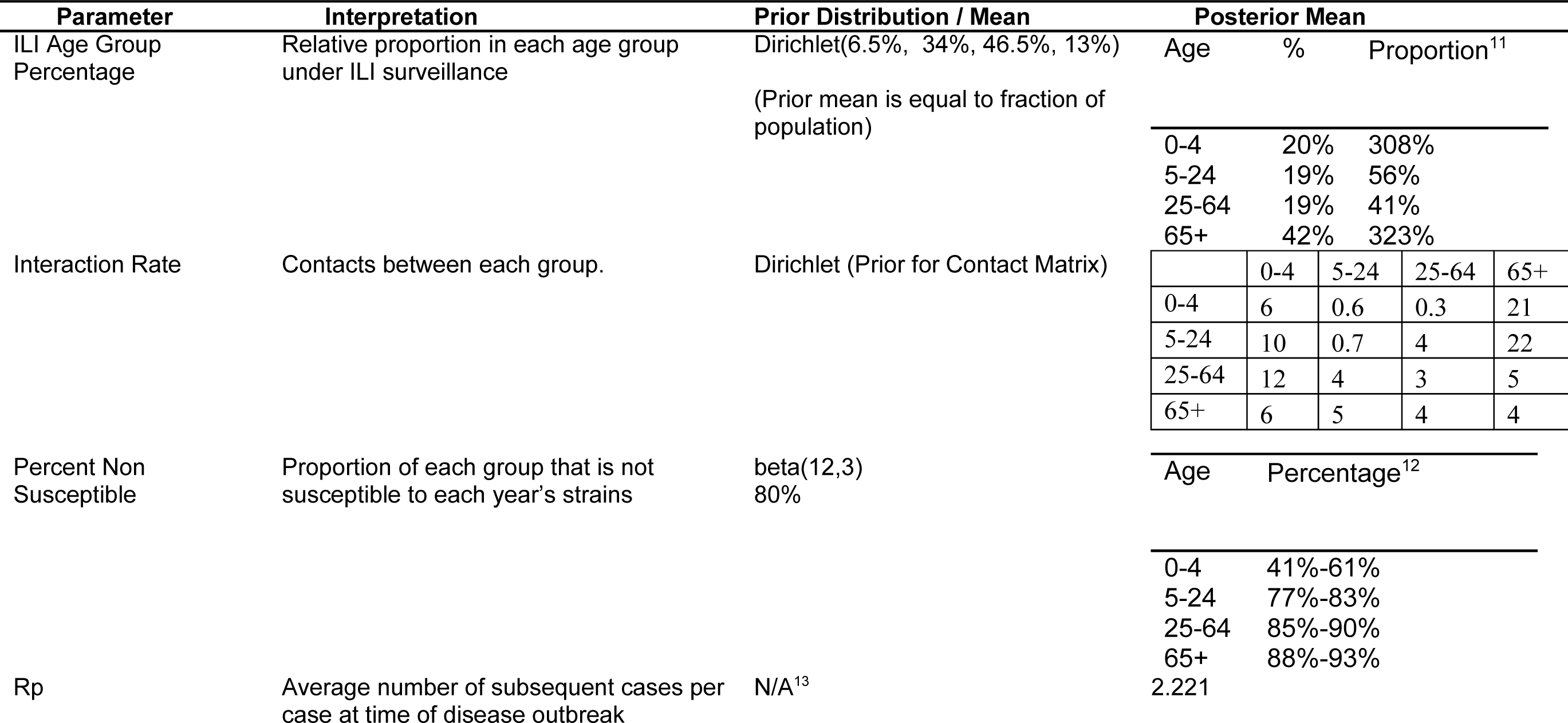
Mean Posterior Estimates

## Model Results

Using historical influenza data for 6 past influenza seasons, 2005 through 2012, excluding 2008 and 2009 due to the outbreak of H1N1, the source data available was used to estimate the various identified model parameters in Table 1. This analysis also provides posterior samples for the joint distribution, including those for the relationships between the number of patients and the reported biosurveillance data. Because of the computational complexity of the model and the limited computation budget for this project, 4 chains with 250 samples after a 50-iteration HMC adaptation period were generated. (This is fewer samples and a significantly smaller adaptation window than would be ideal.)

The output from the model was used to estimate R_p_, used for understanding transmissibility of the influenza virus in the population. Figure 3 shows a histogram of the R_p_ found in the posterior simulations for each year. The estimated ranges are large, showing that the model estimates of transmission dynamics are highly uncertain in the model. In the literature, influenza Rp is estimated as approximately 1.3 for seasonal flu (Chowell et al., 2006), similar to estimates found in a meta-analysis (Biggerstaff et al., 2014), and near the mode of the model estimates^14^.

**Figure 3:**
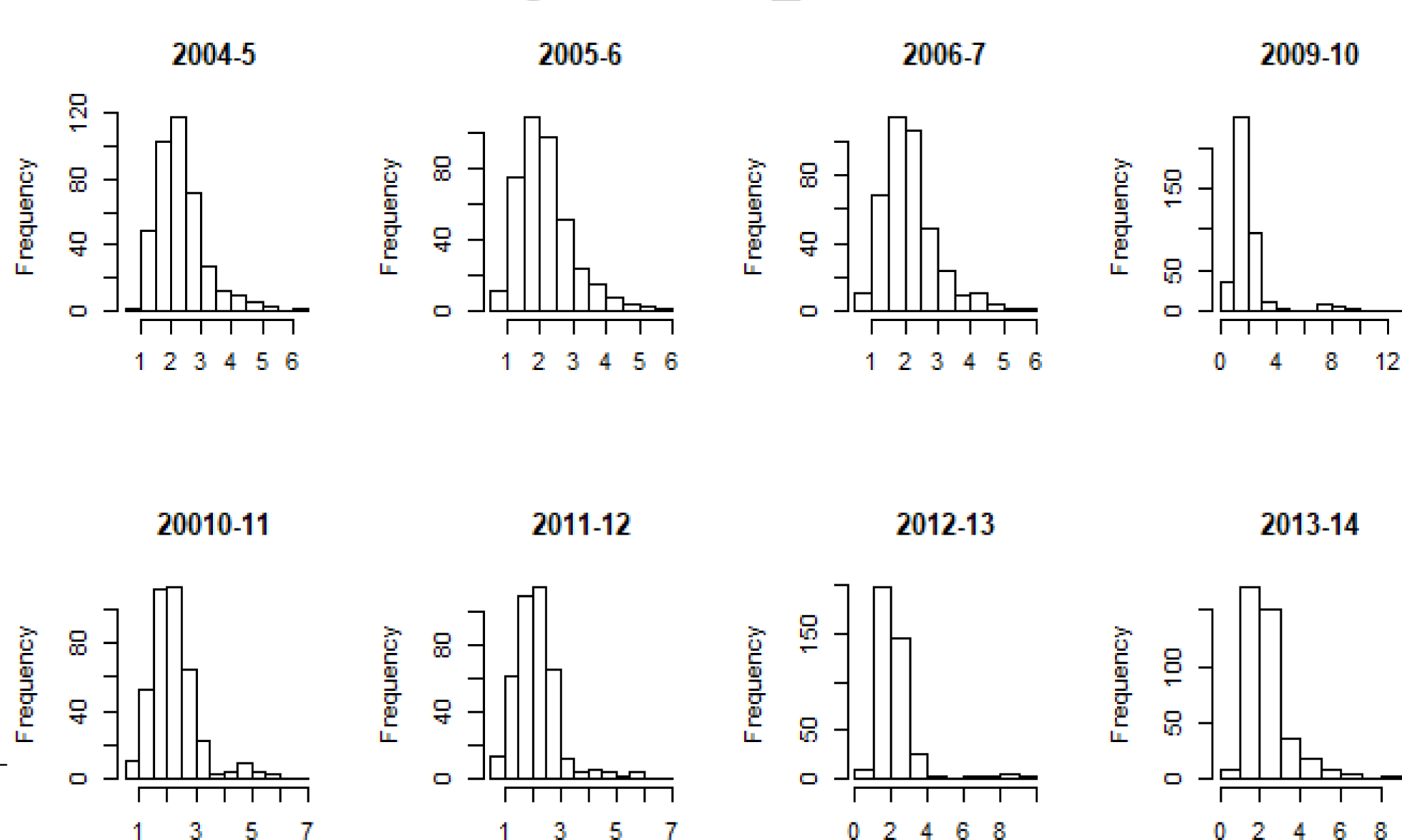
R_p_ Estimates.

## Conclusions

The construction of the model illustrates the feasibility of the approach in general, and wider application of generative models to allow incorporation of novel data sources is a potentially useful general technique. The specific model used is easily extensible. For example, including Google Flu data into the model required two statements for representing the data, another two for specifying the prior distributions, three statements specifying the auxiliary variables needed for characterizing the source and its relationship to the SEIR model, and four lines of code^15^ to specify the relationship between these variables and the rest of the model.

Several limitations for the use of these methods were found in this work, including challenges with non-identifiability when constructing the information model, computational complexity of using Bayesian HMC in tandem with the available differential-equation solvers, and difficulties reparameterizing prior distributions to allow convergence. These are typical challenges when applying the class of Bayesian techniques used, but should all be noted for future applications.

Future work on this model could focus on three significant areas; 1) increased resolution, 2) strain-specific modeling, and 3) improving model convergence and computational efficiency. Already-available disaggregated geographic data on influenza could be used in a multilevel model if the model was extended to allow for it. The information model as developed can easily be extended in this way to use both additional aggregated data sources and disaggregated ones.

This could be done even without a SEIR model that accounts for geographic locations and correlations, which is another useful albeit more computationally expensive extension. The current model using a single strain can further (or alternatively) be extended to represent multiple strains, though the complexity of multi-strain SEIR models grows quickly with the number of strains modeled.

Model convergence could potentially be assisted by improving the parameterization used, which would likely also improve computational efficiency. This requires substantial expertise in the creation and application of these models which the author unfortunately lacks. Model efficiency can alternatively be improved using a set of techniques currently being developed for improving performance of Bayesian inference. The latter includes novel techniques specific to differential equation models, such as the recent work by Margossian, (Margossian, 2017) as well as the ongoing development work on Stan that has led to several significant performance improvements over the past several years. In addition to the direct benefit of improved model performance, such improvements would allow for computationally feasible extension of the model, making the earlier two suggested extensions more feasible.

Overall, this class of model seems to provide a potentially valuable method for rapidly incorporating novel epidemiological data sources into extent models.

## Acknowledgments

Advice for building the models used was provided by several STAN developers. The original ODE model which was used as a template and basis for the model developed here was created by Michael Betancourt, and he and Bob Carpenter provided helpful comments for understanding various issues with convergence of the model. Jim Savage provided excellent advice about the research and model building process, and the process of building the model would have further benefited if that advice had been followed even more carefully. Katherine Laskey provided helpful insight about the issues with identifiability, Steven Popper provided very helpful stylistic and writing guidance, and Paul Davis provided invaluable advice about the research in general, as well as insights on almost every aspect of the work.

## Footnotes

1 The data and material supporting this work can be found in the support files here.

2 The Google Flu data source was discontinued, but functions as a useful example. For decision purposes, it could presumably be gathered again if found to be valuable. It is also a reasonable proxy for several proposed or trialed systems that use live social media or internet data such a Twitter monitoring.

3 This status is usually referred to as “Infected,” based on the terminology of SIR models, where there is only a single compartment for those who have the infection. Here, exposed individuals are also infected, so the compartment is more clearly referred to as infectious.

4 For a basic discussion of the various approaches that exist for modeling disease, including this type of compartmental model, see Chapter 2 of Manheim et al. (Manheim et al., 2016) For a more in-depth look at building compartmental models, the standard textbook, by Anderson and May, is excellent. (Anderson, May and Anderson, 1992)

5 These age groups were chosen to match age groups provided in the CDC ILl data, as well as because it can allow representing school-aged children for understanding possible interventions like school closures, and representing working adults for workplace interventions.

6 The data also reports specific strains of influenza, which a more complex influenza model that represented individual strains might be able to utilize. This capability would also be critical for identification of pandemic strains.

7 The charts representing this data was released in near-realtime during the time the system was in use.

8 For more technical details on this, see Andrew Gelman’s textbook, (Gelman et al., 2014). For details on the various sampling and simulation algorithms used, see Stan^®^documentation (STAN Development Team, 2017) and related technical papers. For an explanation of warm-up for HMC as it occurs in STAN, see the discussion of terminology in comments on Gelman’s blog (Gelman, 2017)

9 Multipliers next to the parameters center the mean so that the Hamiltonian Monte Carlo Sampler is more computationally efficient. (Parameters are log-transformed for sampling, so values close to the bound are hard to sample. This affects gamma distributions near 0, and beta distributions near 0 or near 1.)

10 This value was expected to be significantly higher than the literature suggests for influenza; this is an artifact of accounting for delay in the exposure transition, which ordinary differential equations cannot model directly. Standard methods for accounting for this require more complex differential equations.

11 The relative percentage of ILI participation to actual percentage of population; 3 times as many infants are seen with ILI symptoms as would be expected if all groups were equally likely to have a doctor report their symptoms.

12 The percentage non-susceptible differed between years. Percentages given are the range of the mean values found.

13 The calculation of Rp is not done within the model, and was instead calculated in R based on the model results using the Next Generation Matrix (NGM) method, and means no explicit prior was specified. See Diekmann, Heesterbeek and Roberts for an explanation of the methodology used. (Diekmann, Heesterbeek and Roberts, 2010)

14 In a Bayesian analysis, the mode of the posterior is the maximum a posteriori estimate, which roughly corresponds to a maximum likelihood estimate, and seems to be an appropriate value for comparison.

15 Excluding defining two temporary variables used for transforming the data for the use of array operations, and another used for simplifying the computation of the standard deviation.

